# Universal NicE-seq for high resolution accessible chromatin profiling for formaldehyde fixed and FFPE tissues

**DOI:** 10.1101/2020.04.27.064592

**Authors:** Hang Gyeong Chin, Zhiyi Sun, Udayakumar S. Vishnu, Christina Hao, Paloma Cejas, George Spracklin, Pierre-Olivier Estève, Shuang-yong Xu, Henry W. Long, Sriharsa Pradhan

## Abstract

Accessible chromatin plays a central role in gene expression and chromatin architecture. Current accessible chromatin approaches depend on limited digestion/cutting and pasting adaptors at the accessible DNA, thus requiring additional materials and time for optimization. Universal NicE-seq (UniNicE-seq) is an improved accessible chromatin profiling method that negate the optimization step and is suited to a variety of mammalian cells and tissues. Addition of 5-methyldeoxycytidine triphosphate during accessible chromatin labeling and an on-bead library making step substantially improved the signal to noise ratio while protecting the accessible regions from repeated nicking in cell lines, mouse T cells, mouse kidney, and human frozen tissue sections. We also demonstrate one tube UniNicE-seq for FFPE tissue section for direct NGS library preparation without sonication and DNA purification steps. These refinements allowed reliable mapping of accessible chromatin for high resolution genomic feature studies.

## Introduction

The eukaryotic nuclear genome is packaged into chromatin, consisting primarily of DNA, proteins and RNA, which is further condensed into larger folded chromosome structures during cell division. During cellular events chromatin undergoes remodeling providing accessibility to DNA binding proteins including transcription factors (1–3). Gene promoters and enhancers participate in gene expression and confer to accessible chromatin structure. Recent genome wide methods and studies for mapping chromatin accessibility (open chromatin), nucleosome positioning and transcription factor occupancy have utilized a variety of methods including DNase hypersensitive region sequencing (DNase-seq, 4), Assay for Transposase-Accessible Chromatin using sequencing (ATAC-seq, 5) and nicking enzyme assisted sequencing (NicE-seq, 6). Although both DNase-seq and ATAC-seq are powerful methods, they both require specific reagents and cell type specific optimization including cell number to enzyme/Tn5 transposon concentration and time of incubation (4, 5). While ATAC-seq works primarily on unfixed cells, mitochondrial DNA sequence contamination was a major issue untill a modified Omni-ATAC-seq protocol was developed (7). Our previously published NicE-Seq method also required similar enzyme titration in each cell type to determine an optimal enzyme to cell ratio for effective labeling and capture of accessible chromatin regions. Therefore, all the current methods required a careful titration of cells to enzyme and incubation time for optimal digestion to capture accessible chromatins. Here we report an improved, fast, accurate, and robust method, UniNicE-seq, which generates higher data quality for interrogation of accessible chromatin and eliminating the need for cell number to enzyme titration. We tested this method on a diverse set of cell lines, native and formaldehyde fixed tissue nuclei, fresh frozen formaldehyde fixed 5-10 μM human tissue sections along with FFPE tissue sections and observed markedly improved accessible chromatin signals. Taken together, Universal NicE-seq is a simple, cost effective, nick translation-based method to profile accessible chromatin using NextGen sequencing. Importantly, 5mC incorporated accessible chromatin remain resistant to further degradation eliminating time consuming titration and would allow automation on a variety of biological samples.

## RESULTS

### Universal NicE-seq protects labeled accessible chromatin against enzymatic degradation

During the previously published NicE-seq labeling reaction, the nucleotide mixture contained dNTPs supplemented with biotin-14-dATP and biotin-14-dCTP (6). This resulted in labeling accessible chromatin with biotinylated nucleotides for library preparation and enrichment. However, in the presence of excessive enzyme, the accessible regions were repeatedly nicked resulting in poor library quality and a higher signal to background ratio. In fact, the biochemical property of Nt.CviPII would allow nicking of an unmodified CCD but not mCCD (Fig 1A). To prevent repeated nicking at the same site, we substituted the dCTP in the labelling reaction with of 5-methyldeoxycytidine triphosphate (5-mdCTP). Thus, all available deoxycytidine triphosphates were indeed either 5-mdCTP or biotin-14-dCTP. This modification ensured incorporation of 5-mdCTP and/or biotin-14-dCTP at the nicking sites of Nt.CviPII by DNA Pol I. Modification of CCD sites at the 5’ cytosine renders Nt.CviPII resistant to further nicking. The second modification step we implemented was on-bead library preparation which helped in reducing the background and enhancing the signal to noise ratio for the accessible region of the genome. For the UniNicE-seq, after labeling reaction the biotin labeled DNA of HCT116 cell was isolated, sonicated, and captured on streptavidin magnetic beads for library preparation and sequencing (Fig 1B). A slight modification to this protocol by substituting dATP to Texas-Red-dATP allowed both visualization and sequencing (termed as NicE-viewSeq) of the accessible chromatin region (Fig 1C, Please contact the authors for step wise protocol).

**Figure 1:**
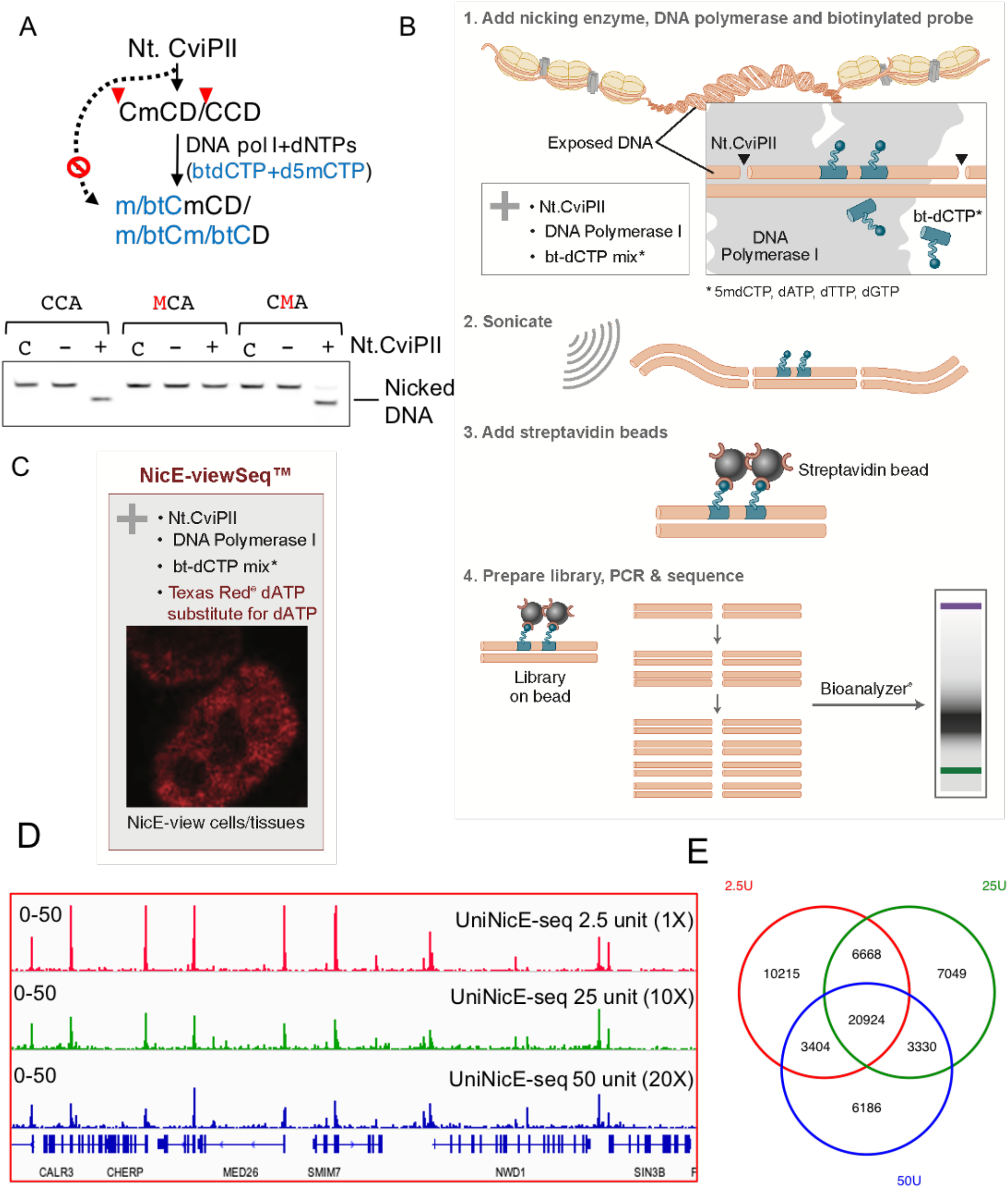
Addition of 5mdCTP in UniNicE-seq work flow. (A) Schematic diagram showing Nt.CviPII is blocked by 5-methylcytosine in recognition (CCD) sequence. During nick translation 5mdCTP can be incorporated at one or both cytosine position. Nucleotide mixtures containing biotinylated dCTP and 5mdCTP would allow both labels at CCD sites. Lower panel shows blocking of nicking by the presence of 5mC at CCD sites. After the nicking reaction the denatured oligonucleotides were resolved in a urea-acrylamide denaturing gel by electrophoresis. C is control input oligos, − or + indicate absence or presence of Nt.CviPII in the reaction. mC and btC represents 5-methylcytosine and biotinylated-cytosine respectively. Nicked oligonucleotide migrates faster on the denaturing urea gel. (B) Schematic diagram of UniNicE-seq method for accessible chromatin library preparation. (C) Substitution of dATP by Texas Red conjugated dATP will allow accessible chromatin visualization in the nucleus. (D) IGV screen shot of the normalized read density of UniNicE-seq using 2.5U (top track), 25U (middle track) and 50U (bottom track) of Nt.CviPII in HCT116 (E) Venn diagram showing overlap of peaks in various amounts of Nt.CviPII in the universal NicE-seq (UniNicE-seq) reaction in HCT116 cells.

To determine the effect of 5mC incorporation in accessible region for library preparation, we performed UniNicE-seq in the presence of 10X and 20X (25 and 50 units) excess Nt. CviPII nicking enzyme. Indeed, neither of the two reaction conditions showed loss of accessible chromatin in HCT116 cells (FRiP 0.19-0.14) demonstrating the robustness of 5mdC labeled accessible chromatin protection against degradation (Supp table 1). In a similar reaction condition using 25 and 50 units Nt.CviPII and original NicE-seq protocol we obtained FRiP score below 0.01 and loss of accessible chromatin peaks (Supp table 1). In addition, 2/3rd of peaks overlapped between universal NicE-seq libraries made with 1X, 10X and 20X nicking enzymes and signal to noise ratio were comparable (Fig 1D and E). These results unequivocally demonstrate the versatility of 5-mdCTP in the reaction to protect accessible regions without titration of cell to enzyme concentration.

To assess other improvements in our new method, we compared the number of accessible chromatin peaks detected and peak height which measures as the ratio of reads within the peak versus background. First, we compared peak numbers between libraries made on beads (on bead) and control (off beads) either with 5-mdCTP or dCTP in the dNTP mix (Supp Fig 1A). The majority of accessible chromatin peaks are common across these libraries as shown in the Venn diagram suggesting that our modifications are not changing the distribution of identified regions (Fig. 2A). Furthermore, the signal to noise ratio was improved in libraries made on beads as observed by peak height and distribution of fold change (FC) values (Fig. 2B). For comparison between original NicE-seq (off bead C) and universal NicE-seq (on bead 5mC) the peak fold changes were better (6 vs. 8) and fraction of reads in the peaks (FRiP) were 0.095 and 0.26 respectively (Supp fig 1B).

**Figure 2:**
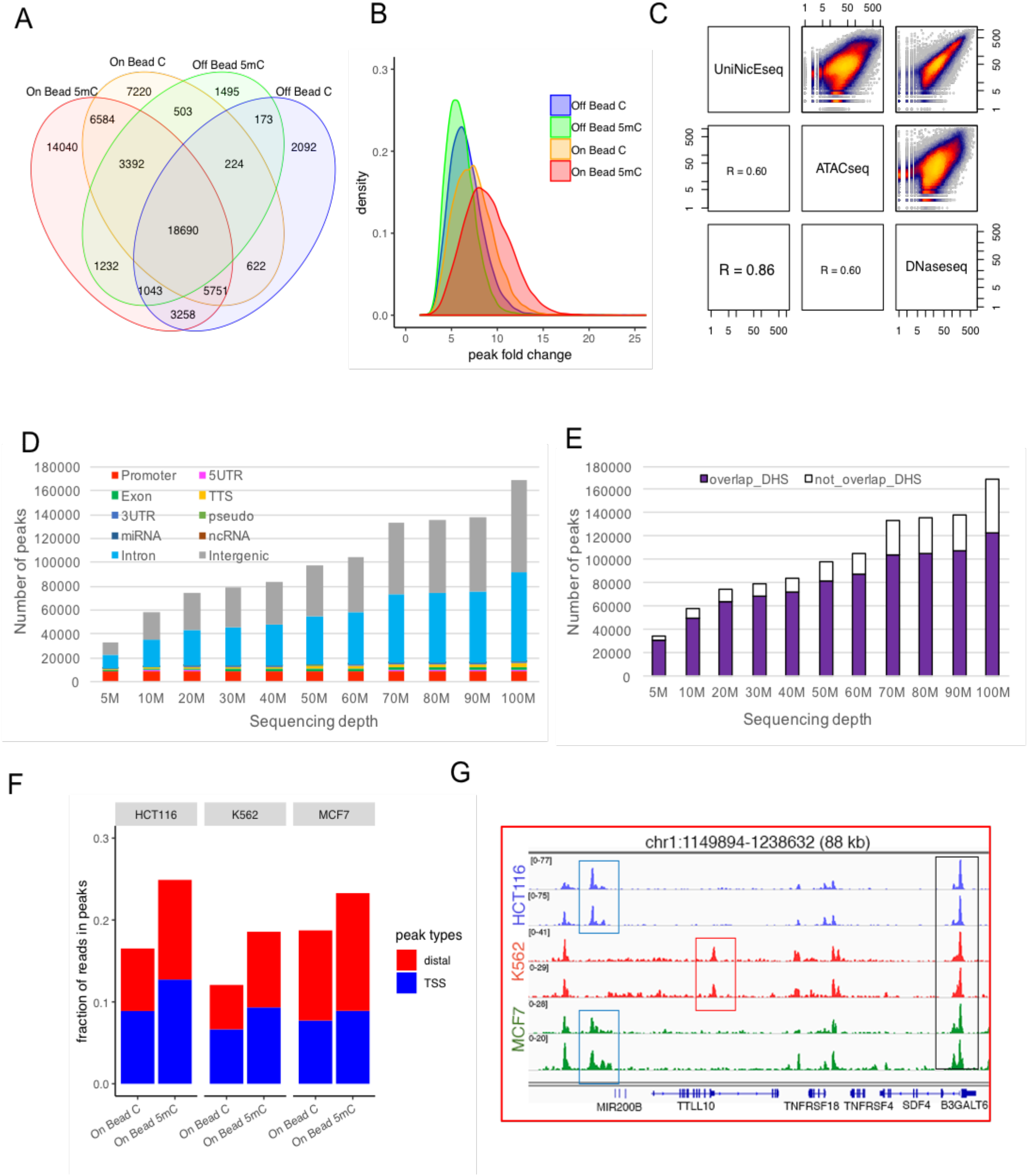
UniNicE-seq optimization and validation. (A) reaction conditions and library making method comparison. Venn diagram showing overlap of peaks in various reaction conditions and library making method. (B) Distribution of fold change (FC) values (derived from MACS2) of the accessible chromatin peaks detected by on bead 5mC, on bead C, off bead 5mC and off bead C methods. Accessible chromatin peaks detected by on bead 5mC (red) showed higher FC values on average than other methods. (C) Pearson correlation of accessible chromatin peak read densities between UniNicE-seq, ATAC-seq and DNase-seq of HCT116 cells. (D) Distribution of HCT116 UniNicE-seq (On Bead 5mC) peaks in different genomic regions at sequencing depth from 5M to 100M alignment pairs. (E) Numbers of HCT116 UniNicE-seq accessible chromatin peaks that overlap and not overlap with the reference human DNase I hypersensitivity sites at sequencing depth from 5M to 100M alignment pairs. (F) Fraction of reads in peaks that map to TSSs (+/−500bp of TSS) and distal elements (>500bp from TSS) from libraries generated using the on-bead UniNicE-seq methods on three human cell lines (HCT116, K562, MCF7). (G) Representative IGV screen shot of the normalized read density of the UniNicE-seq libraries of the three human cell lines, HCT116, K562 and MCF7, from experimental replicates.

We further compared the accessible chromatin reaction conditions and reads between UniNicE-seq, ATAC-seq and DNase-seq of HCT116 using Pearson correlation of peak read densities for data quality evaluation. In terms of data quality, UniNicE-seq and DNase-seq had the highest correlation (R=0.86) compared to UniNicE-seq with ATAC-seq (R=0.60) or ATAC-seq with DNase-seq (R=0.60), demonstrating most DHS (DNase Hypersensitive Sites) are indeed captured in this modified protocol (Fig. 2C). Since nucleosome-free DNA regions differentially affect distant communication in chromatin and accessible chromatins are a hallmark of active gene promoters, we compared accessible chromatin profile and transcription start sites (TSS) in duplicates datasets, across both off-bead and on-bead methods. Indeed, all the TSS were enriched with accessible chromatin tag densities (8). HCT116 cell nuclei labeling in the presence of 5mdCTP displayed highest tag densities and higher signal to noise ratio with high degree of correlation ~99% between technical duplicates (Supp Fig. 2A-C). Comparison between UniNicE-seq, ATAC-seq and DNase-seq peaks of HCT116 showed stronger peak signals in UniNicE-seq sequencing tracts that were derived from equal numbers of unique sequencing reads (Supp Fig 2D), with a high percentage of common accessible regions between UniNicE-seq and DNase-seq (Supp Fig. 2E). In order to define the accessible chromatin coverage of UniNicE-seq we also performed sequencing depth analysis and discovered accessible chromatin regions in promoters and other genic regions. Indeed, at a sequencing depth of 5 million reads we were able to identify most of the HCT116 promoters. At higher sequencing depth of 100 million reads, intronic and intergenic accessible regions were more prominently detectable, although the significance of these regions remains unclear (Fig. 2D). Since UniNiCE-seq and DNase-seq showed a high degree of correlation, we compared both datasets at a sequencing depth of 100 million reads with 10 million read increments and approximately 71% of the UniNicE-seq peaks overlapped with DHS, suggesting strong confidence in the UniNicE-seq method (Fig. 2E). To establish UniNicE-seq as a general protocol, we assayed two additional cell lines, MCF7 and K562, and compared accessible chromatin between libraries made on beads with either with 5-mdCTP or dCTP in the dNTP mix (Supp. table 2). We calculated fraction of reads in peaks (FRiP) analysis of all the peaks generated by UniNiCE-seq to compare with HCT116 cells. The FRiP score was comparable amongst all three cell lines in both the total number of peaks and promoter peaks, suggesting substitution of 5mC for C and on bead library preparation improves the number of accessible chromatin peaks with identical sequencing depth of 11 million pair reads (Fig 2F). As expected, the cell line specific unique and common peaks were also identified in UniNicE-seq (Fig. 2G, Supp Fig 3) with consistently higher numbers of identified TSS peaks (Supp table 2). Taken together, these results demonstrate universal NicE-seq is a superior and robust method than its predecessor NicE-seq protocol to profile accessible chromatin negating the need for enzyme titration and protecting accessible regions for NextGen sequencing.

### Universal NicE-seq of mouse tissue

We further applied the UniNicE-seq to mouse T cells and kidney tissues. T cells were formaldehyde fixed and serial diluted from 25,000 to 500 cells for UniNicE-seq labeling, and libraries were made in duplicate. The signal to noise ratio of all samples were consistent, and MACS2 peak calling was made with down-sized 13M unique non-mitochondrial reads (Supp fig. 4). As expected, the majority of the accessible chromatin peaks were common in all replicates (Fig. 3A). The pairwise total read densities comparison between 500, 5K and 25K cells also displayed strong correlation of r value between 0.95-1. (500 vs. 5K, r=1; 5K vs. 25K, r=0.97; 500 vs. 25K, r=0.95). Since accessible DNA quantity varied between different cell numbers, we further tested if the read density between the common peaks are similar. For validation, average log2 (normalized reads) in 6062 peaks that were detected in all 6 samples were plotted on a 3-dimensional scatter plot. The observation that all data points line up on a straight line in the 3-dimensional space shows that signal in 25000, 5000 and 500 T-cells are highly correlated and that there was no loss of signal with decreased numbers of cells (Fig. 3B). A correlation matrix of Pearson values corroborates the above observation (Fig. 3B). A strong correlation was also observed when all reads were compared (Supp fig 4B). In addition to mouse T cells, we also interrogated and compared accessible chromatin data from formaldehyde fixed with unfixed mouse kidney cells. Accessible chromatin from 25K fixed cells were compared with 25K, 10K, 1K, 500 and 250 unfixed cells. The quality control metrics of UniNicE-seq libraries applied to mouse kidney tissues was good (Supp table 3). The peak numbers between fixed and unfixed cells were similar along with the signal to noise ratio between 25000-500 cells (Fig. 3C, Supp. Table 3). We also observed that the nonfixed high number kidney nuclei (>1K) would form clumps and make the NicE-seq labeling reaction inefficient resulting in decrease in FRiP score (supp. Table 3). Direct comparison between common accessible region peaks of 25000 fixed and non-fixed cells displayed good correlation (P=0.98), suggesting both samples are compatible with the UniNicE-seq protocol, and is robust without the enzyme titration requirement for low cell numbers (Fig. 3D). Heatmap analyses showing normalized RPKM at TSS and enhancers of 24,920 mouse Ref-seq genes and the surrounding 2kb region displayed the central accessible region. Further comparison with mouse kidney H3K4me3, H3K27Ac, PolII and CTCF ChIP-seq data demonstrated an enrichment of accessible regions as expected (Fig 3E). We further compared data sets from the UniNicE-seq method generated from 25K fixed mouse kidney cells and 0.25K non-fixed cells with Omni-ATAC-seq (improved ATAC-seq, 7) and DNase-seq datasets. In experiments involving UniNicE-seq and unfixed 500 cells and Omni-ATAC-seq and 50K cells, 65% of the peaks from UniNicE-seq were common with Omni-ATAC-seq. (Supp fig 5A). Analysis of tag density and peak width as well as FRiP score comparison suggested that UniNicE-seq on 500 unfixed mouse kidney cells yielded high quality accessible chromatin data (Supp fig 5B, C). TSS, PolII and active enhancer and chromatin marks were also prominent with 500 kidney cells (Supp fig 5D, E) These results show that genes with active promoters and enhancers are captured by UniNicE-seq as open chromatin regions, regardless of fixation and with small cell number from tissue samples.

**Figure 3:**
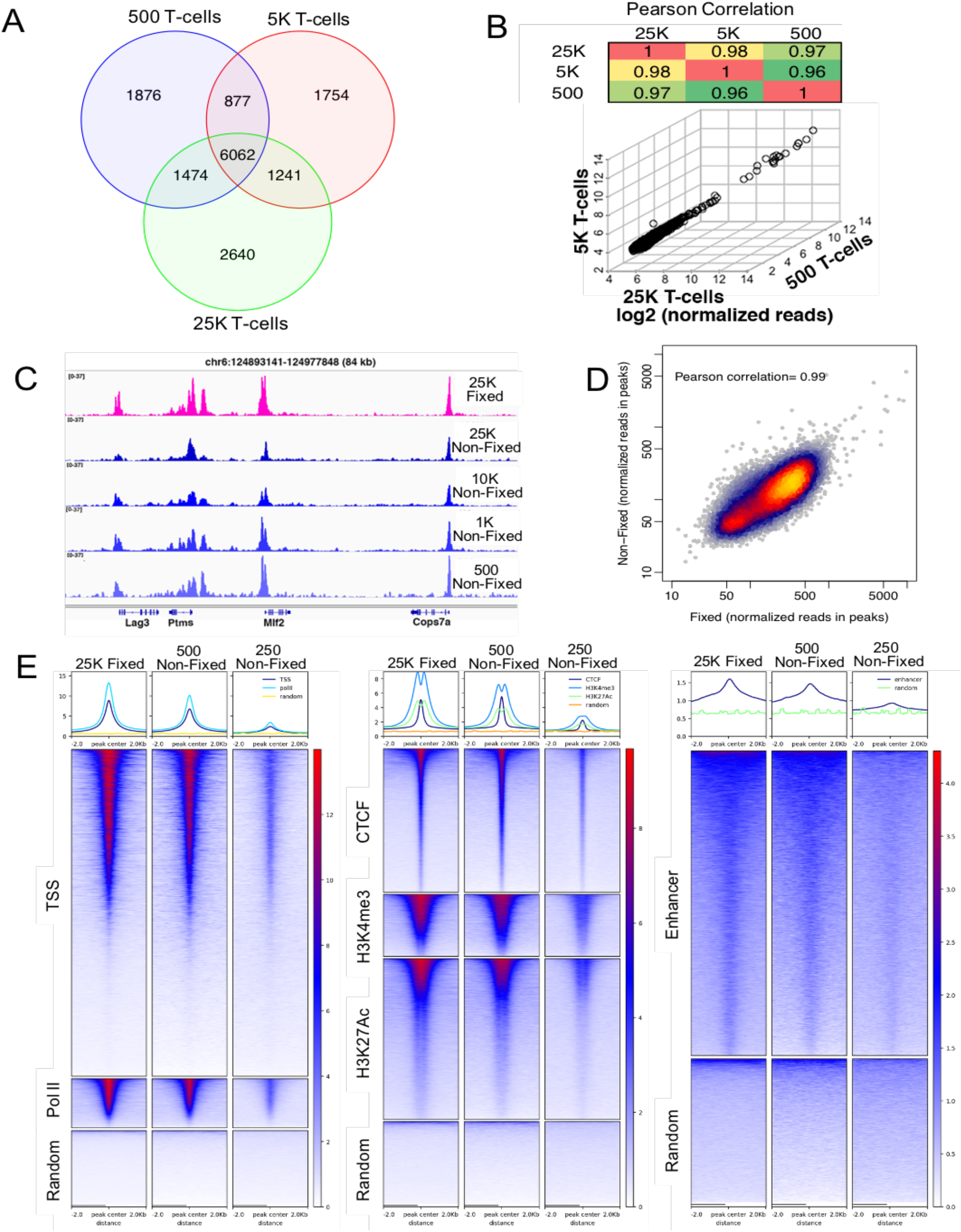
UniNicE-seq of mouse T cells and kidney tissue. (A) Venn diagram demonstrating common accessible regions in mouse T cells with sample size from 25K, 5K and 0.5K cells. (B) 3-dimensional scatter plot of technical duplicate T-cell samples from 25K, 5K and 0.5K cells. Average log2 (normalized reads) in common accessible chromatin peaks between all samples are plotted. A correlation matrix of Pearson values of the common accessible chromatin peaks is shown on top. (C) Representative IGV screen shot of the normalized read density of the UniNicE-seq libraries of mouse kidney nuclei. Comparison between formaldehyde fixed nuclei and non-fixed nuclei is shown. Non-fixed nuclei numbers were between 100K to 0.25K. (D) UniNicE-seq libraries of 25K fixed and non-fixed mouse kidney cells were downsized to 21.5M properly aligned read pairs (after removing mitochondrial reads and PCR duplicates). MACS2 called 40716 narrow peaks from the fixed sample and 27853 peaks from the non-fixed sample, among them 15901 are common to both samples. The read density of the 2 libraries in the set of common peaks are highly correlated with a Pearson correlation efficiency of 0.99. (E) Heatmap showing normalized RPKM of 100K fixed, 0.5K nonfixed and 0.25 K non-fixed mouse kidney nuclei at TSS, PolII and random along with chromatin features including CTCF, H3K4me3, H3K27Ac and random fragments.

### Universal NicE-seq of human 5-10 μM frozen tissue sections

Clinical samples from patients are often limited and increasingly obtained from fresh biopsies. Routinely, 5-10 μM tissue sections are used for immunopathological and other staining analysis. Currently, ATAC-seq method for chromatin accessibility studies requires 50,000 purified nuclei from patient brain and cartilage tissue samples (7, 9). Indeed, 20 mgs of brain tissue samples were used from post-mortem human brain samples for nuclei preparation (7). However, obtaining large tissue volume from live patients is inconvinient, and often donor samples are limited. Therefore, we attempted UniNicE-seq using single frozen 5-10 μM tissue sections, mimicking biopsy samples. The protocol was slightly modified for this application, with a formaldehyde fixation step of the tissue section prior to the labeling reaction on the slide. The fixing ensured the attachment of tissue section for on-slide labeling reaction and subsequent washing steps. We than applied the UniNicE-seq protocol to make accessible chromatin libraries from tissue sections. Accessible chromatin regions were revealed, as expected, between two different lung tissue sections (Supp Fig 6A, Supp table 4). Total read counts between these two tissue sections displayed strong correlation r=0.97 (Supp Fig 6B). Similarly, all common accessible peak reads between both tissue sections also were highly correlated (r=0.97) demonstrating high quality of common accessible regions (Supp Fig. 6C).

Following the reproducibility of accessible chromatin of lung tissue, we then applied the UniNicE-seq protocol to make accessible chromatin libraries from human adult and fetal tissue including liver, lung and kidney, pooled both rep[licates and down sized the reads and compared accessible chromatin landscape amongst them. Accessible chromatin regions were revealed, as expected, across all tissue sections (Fig 4A). Active enhancer and promoter histone marks, particularly H3K4me1, H3K27Ac, H3K4me3 coincided with all accessible regions identified by UniNicE-seq and correlated with curated promoters in human adult lung tissue compared to fetal lung tissue (Fig. 4B). Across all tissues accessible chromatin was enriched in promoter, 5’UTR, and rRNA genic regions demonstrating commonality further supporting our method is compatible with the identification of open chromatin peaks in tissue sections (Fig 4C).

**Figure 4:**
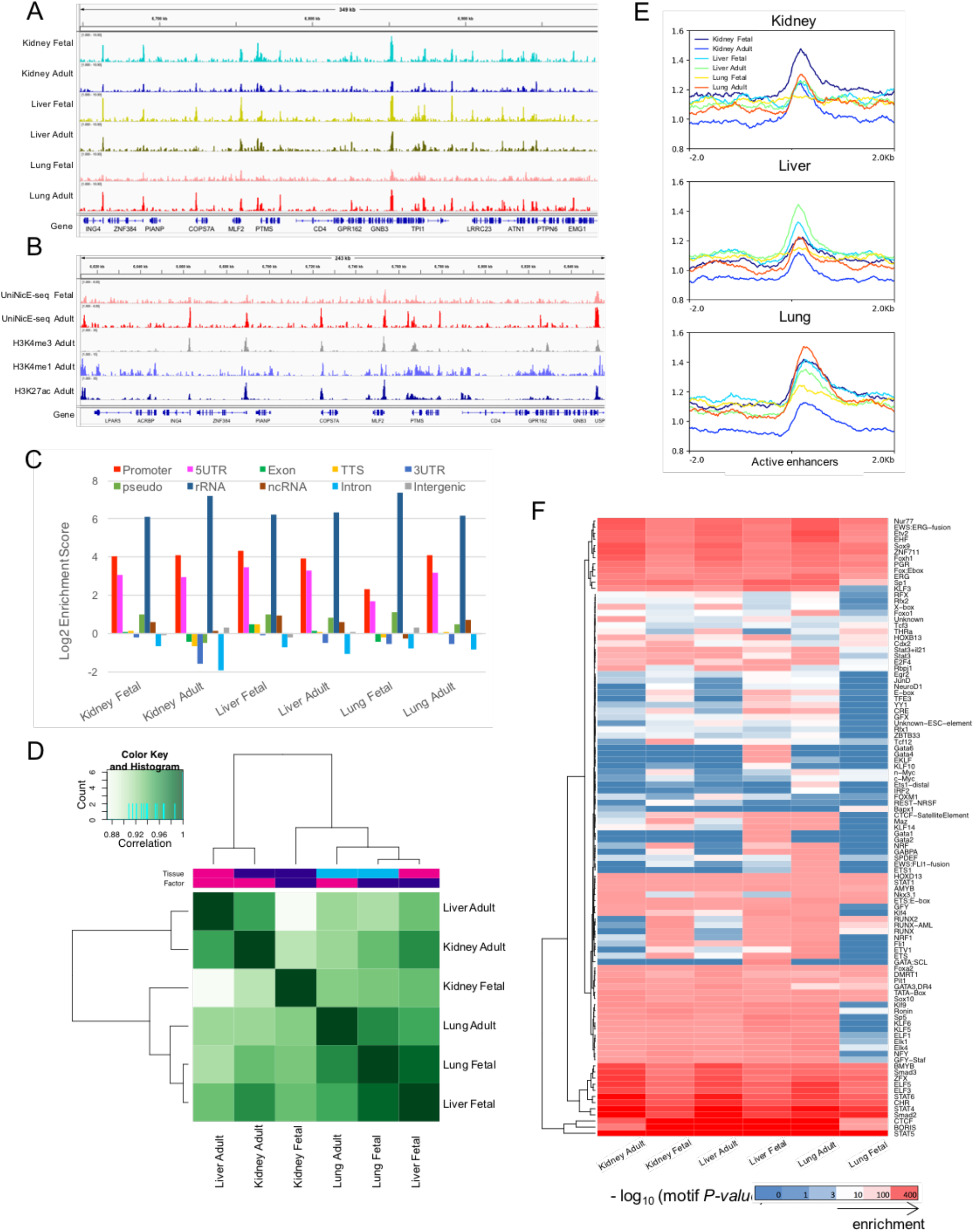
UniNicE-seq of 5-10 μM human frozen tissue sections and genomic features analysis. (A) Representative IGV screen shot of the normalized read density of the UniNicE-seq libraries of human fetal and adult tissue sections derived from kidney, liver and lung. (B) Representative IGV screen shot of the normalized read density of the UniNicE-seq libraries of fetal and adult lung tissue sections with ChIP-seq chromatin marks of adult tissue for H3K4me3, H3K4me1 and H3K27Ac. (C) Enrichment of UniNicE-seq peaks with various genomic features. (D) Correlation analysis of UniNicE-seq peaks of human fetal and adult tissues. (E) Profile plot of UniNicE-seq open-chromatin signals (peak density) of human tissues samples in Kidney, liver and lung tissue-specific active enhancers (AE). (F) Heatmap representing the enrichment of consensus TF binding motifs identified in each tissue sample near the UniNicE-seq peaks. Both of the TF binding motifs and the samples are organized by unsupervised k-means clustering method. The p values of e-6 was considered for the cluster analysis.

Cells undergo transcriptional changes during differentiation, especially the acquisition of tissue specific enhancers. Therefore, we profiled enhancers in fetal and adult lung tissues using known histone marks associated with enhancers. However, in fetal and adult lung there was a marked difference in accessibility of promoter and 5’UTR regions. When we compared adult vs. fetal tissues a varying degree of correlation was observed among all 3 different tissue types (Fig 4D). Accessibility of active enhancer elements in kidney was more enriched in fetal tissue compared to liver enhancers that predominated in the adult tissue. As expected, active enhancers in adult lung tissues were more enriched compared to fetal tissue suggesting a developmental cue for enhancer activation post birth (Fig 4E). Similarly, transcription factor consensus binding site near the UniNicE-seq derived accessible chromatin binding region displayed varying degrees of similarity and a contrast between fetal and adult tissues, with fetal tissue demonstrating developmental and functional programming of lung tissue (Fig 4F). Principal component analysis of read counts between fetal and adult tissue accessible chromatin displayed close similarity in liver compared to kidney and lung tissue (Supp Fig 6D). However, transcription start sites were always accessible in both fetal and adult tissue to varying degrees (Supp Fig 6E).

### One-tube universal NicE-seq protocol of human 5-10 μM FFPE tissue sections

Change in chromatin accessibility underlie various diseases including cancers and are often available as FFPE samples. Therefore, we modified universal NicE-seq to be compatible with FFPE tissue sections. The major goal was to develop the reaction protocol that will negate DNA purification and sonication step and will directly feed into NGS library preparation (Fig 5A).

**Figure 5:**
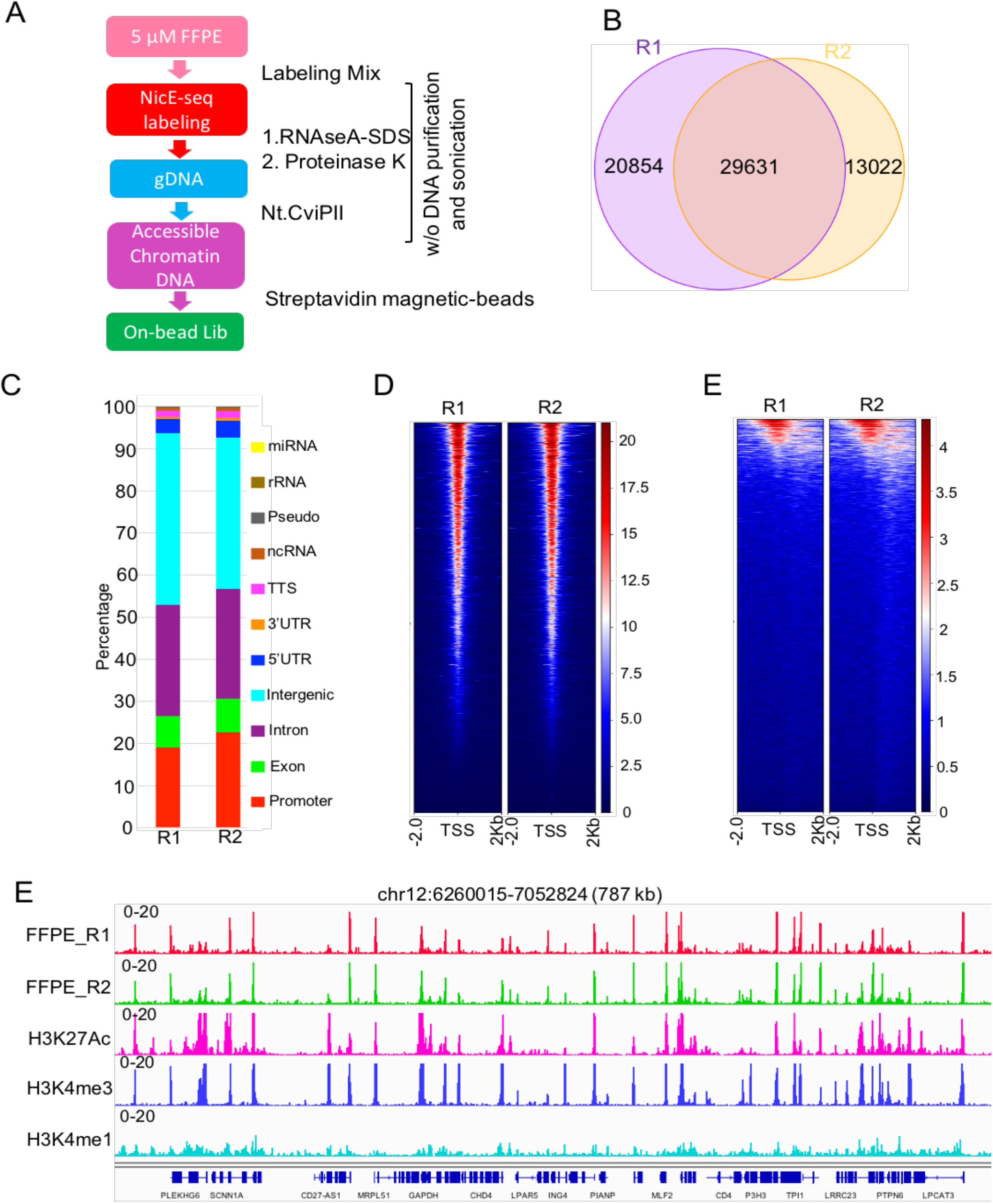
One tube UniNicE-seq of 5-10 μM human liver FFPE tissue sections. (A) Work flow diagram of one tube universal NicE-seq (B) Venn diagram demonstrating common accessible regions in two different human 5-10 μM liver normal FFPE tissue sections designated as R1 and R2. (C) Peak annotation of accessible peaks of liver FFPE tissue section showing the genomic feature distribution. (D) Heatmap and signal intensity profile plot of TSS (that includes ± 2Kb of flanking region) in liver FFPE tissue section. (E) Heatmap and signal intensity profile plot of active enhancer elements (that includes ± 2Kb of flanking region) in liver FFPE tissue section. (F) Representative IGV screen shot of accessible chromatin peaks of two liver FFPE tissue section and histone marks, H3K27Ac, H3K4me3 and H3K4me1.

We introduced a few notable changes. For removal of paraffin we introduced warm paraffin oil wash in place of organic solvent, which is flammable and would need special handling. After paraffin oil based removal of paraffin, the tissue sections were hydrated and equilibrated in PBS for compartibility of labeling reaction. Once the labeling reaction was over, RNaseA was added and crosslinking was reversed at 65°C incubation. The reaction mix was treated with proteinase K for removal of any protein to enrich DNA in the tube. In the next step proteinase K was heat inactivated and nicking enzyme Nt.CviPII was added again to the reaction. Since Nt.CviPII is sensitive to newly labeled accessible chromatin incorporated with 5mC, it would digest away all other DNA of the geome thus enriching only accessible regions. These accessible DNA fragments were captured on strpavidine beads for on-bead NGS library preparation and sequencing (Please contact the authors for step wise protocol).

To demonstate the proof of principle, two different human liver FFPE samples were profiled for accessible chromatin studies (Supp table 5). Both tissue sections displayed 60-70% common accessible regions and a strong correlation of read counts (Fig 5B, Supp fig. 7). Most accessible regions were enriched in promoter, exon, intron and intragenic regions as expected (Fig 5C, D, E). IGV browser display showed accessible chromatin peak correlation with active chromatin marks, H3K27ac and H3K4me3 along with active enhancer mark, H3K4me1 (Fig 4F). Therefore, we concluded that universal NicE-seq works well in single FFPE tissue section.

## Discussion

In summary, we demonstrated the ease of application of UniNicE-seq to a variety of human cell lines, mouse tissues and human organ derived tissue section from both frozen-fixed and FFPE samples for obtaining chromatin accessibility information. Two protocol modifications, 5mdCTP during labeling and an on-bead capture of labeled DNA and library preparation led to enhance robustness without the need for significant optimization. This improved UniNicE-seq method employs a single protocol without any enzyme tittering for optimization of the reaction time to enable accessible chromatin mapping in both unfixed and formaldehyde fixed cells, and 5-10 μM frozen tissue sections in contrast with Tn5 transposon-based method where the transposon simultaneously fragment and tag the unprotected regions of DNA and often needs labor intensive optimization. Furthermore, we have demonstrated that universal NicE-seq is compatible with FFPE tissue sections without DNA purification and sonication. This is particulary important since small cell numbers or limited biological material do not yield large amounts of DNA, and any purification step would result in loss of the DNA. Similarly, sonication mediated DNA sizing on limited amounts of DNA is not advisable. Therefore, one-tube universal NicE-seq will simplify the workload and be more consistant.

Till date, ATAC-seq has not been shown to work on FFPE tissue section. However, it may be noted that integrity of DNA is essential for any accessible chromatin mapping in FFPE tissue samples, since most accessible regions are ~ 500 bp length. Thus, UniNicE-seq offers one step “labeling and protection” of accessible chromatin. Furthermore, UniNicE-seq of 5-10 μM frozen tissue and FFPE sections doesn’t need nuclei purification, thus have the potential to parallel process multiple tissue sections. Indeed, in a recent pan cancer accessible chromatin studies using ATAC-seq the authors have used 20 mgs of cancer tissues to isolate nuclei for library preparation (10). These features demonstrate that UniNicE-seq is a robust method for use with frozen and fixed developmental tissue samples that could be extended to include clinical tissue samples which are currently difficult to obtain in large quantity for chromatin accessibility studies.

## Methods

### Cell culture

HCT116 (ATCC, USA) cells were cultured in McCoy’s 5A media supplemented with 10% fetal bovine serum. MCF7 and HeLa cells (ATCC, USA) were cultured in DMEM plus 10% FBS.

#### Accessible or open chromatin labeling of cells and tissue sections

25,000 cells HCT116 cells were used for routine library construction. Cells were cross-linked using 1% formaldehyde for 10 minutes at room temperature and quenched by using 125 mM glycine. Nuclei were isolated by incubating the cross-linked cells in cytosolic buffer (15 mM Tris-HCl pH 7.5, 5 mM MgCl_2_, 60 mM KCl, 0.5 mM DTT, 15 mM NaCl, 300 mM sucrose and 1% NP40) for 10 minutes on ice with occasional agitation. Nuclei were precipitated by spinning at 1000 × *g*, 4°C for 5 minutes and supernatant were discarded. The nuclei pellet was resuspended in 100 μls of 1x PBS. Open chromatin DNA was labeled with biotin by incubating the nuclei in presence of 2.5 U of Nt.CviPII (NEB, R0626S), 50 U of DNA polymerase I (NEB, M0209S) and 30 μM of each dGTP and dTTP, 24 μM of 5mdCTP (NEB, NO356S) and dATP 24 μM plus 6 μM of biotin-14-dATP (Invitrogen, 19524016) and 6 μM of biotin-14-dCTP (Invitrogen, 19518018) at 37°C for 2 hrs with occasional mixing. The labeling reaction was stopped with 20 μL of 0.5 M EDTA, and 2 μg of RNase A was added to the labeling reaction and incubated at 37°C for 0.5 hour to digest RNA. The DNA was isolated using phenol-chloroform extraction and ethanol precipitation method or Qiagen spin column (Qiagen, USA). For NicE-seq libraries using 25 or 50 units enzyme the reaction conditions were identical except that the incubation was at 800 RPM at 37°C for maximal NicE-seq labeling.

For tissue samples, mouse T cells, human liver and mouse kidney cells, nuclei were prepared, formaldehyde fixed before labeling reaction, as described above. In experiments involving unfixed tissue nuclei, the nuclei suspension was in 1X PBS before labeling.

Frozen tissue sections of lungs on slides were procured from US Biomax, Inc. (HuFTS251). For frozen 5-10 μM tissue section fixing and labeling, the slides were treated with 1% formaldehyde for 10 minutes at room temperature and quenched by using 125 mM glycine. After 1xPBS wash, the slide was treated with cytosolic buffer (15 mM Tris-HCl pH 7.5, 5 mM MgCl_2_, 60 mM KCl, 0.5 mM DTT, 15 mM NaCl, 300 mM sucrose and 1% NP40) for 10 minutes at 4°C. 200 μl of the labeling buffer was placed on the tissue section and the slide was transferred to a humidified chamber at 37°C for 2 hrs. (Please contact the authors for step wise protocol).

#### Accessible or open chromatin labeling of FFPE sections

FFPE slides of the liver sample were custom ordered from BioChain, USA, with 24-48 hrs formalin fixation. Slides were mounted with 5μM sections. The slide were incubated at 52°C for 20 min with mineral oil (Sigma, USA), followed by gradual ethanol wash (100%, 90%, 80%, 70% ethanol) in jars at room temp for 5 min each step. The final wash was in milliQ water for 2 min. Then, the slides were incubated in PBST buffer (1xPBS in 0.1% Tween 20) at 65 °C for 1 hr, and incubated again in 1X PBST buffer containing protease inhibitor (Sigma, USA) at room temperature for 10 min, washed by 1X PBS buffer, and dried at the room temperature. The slides were rehydrated with 1X PBS buffer and treated with the cytosolic buffer for 10 min at 4°C. Cytosolic buffer was washed away and the NicE-seq labeling reaction was identical to the frozen tissue section procedure.

After the labeling reactions, 20 μls (200 units) of proteinase K (NEB, USA) and 20 μl of 20% SDS was added and the tube was incubated at 65 °C for o/n. Proteinase K was inactivated by incubating the reaction tube at 95 °C for 2 min. We next added 2.5 units of Nt.CviPII to the reaction mix and incubated for 16 hrs at 37 °C. Nt. CviPII enzyme in the reaction was heat inactivated at 65 °C for 15 min. These processes ensured that accessible chromatin fragments in the solution for strepavidine magnetic bead enrichment as described below. Please refer to supplementary text1: stepwise detailed protocol: universal NicE-seq (25000-250000 HCT116 cells), Appendix 4.

#### Selective enrichment of labeled accessible/open chromatin

The isolated genomic DNAs (~200 ngs/25K cell) were sonicated into 150 bp fragments (Covaris) and the entire reaction product was mixed with 20 μL of Streptavidin magnetic beads (Invitrogen 65001, blocked using 0.1% cold fish gelatin in 1 × PBS overnight at 4°C) in 1 mL of B&W buffer (10 mM Tris-HCl pH 8.0, 1 mM EDTA, 2 M NaCl). Biotin-labeled open chromatin DNA was captured by streptavidin at 4°C for 2 hours with end-over-end rotation. The beads were washed four times with B&W buffer plus 0.05% of Triton X-100 followed by one-time wash with TE. The beads were resuspended in 50 μL of TE. The DNA was end-repaired, washed twice with B&W buffer plus 0.05% of Triton X-100, dA-tailed and washed with B&W buffer plus 0.05% of Triton X-100. And finally, NEB Illumina adaptor (NEB, E7370S) was ligated and washed twice with B&W buffer plus 0.05% of Triton X-100. A final wash of the bead bound DNA was performed with TE and the bound DNA was resuspended with 20 μls of TE. 10-20 μL of bound DNA was used for library amplification using PCR (NEB, E7370S). Routinely 8-10 PCR cycles were used to generate enough amount of library DNA for sequencing. Low input cell numbers between 250-500, library was amplified 12-13 cycles. The library was examined and quantitated with high-sensitive DNA chip (Agilent, 5067-4627).

### Bioinformatics analyses

#### Data processing and peak calling

Adaptor and low-quality sequences were trimmed from paired-end sequencing reads using Trim Galore (http://www.bioinformatics.babraham.ac.uk/projects/trim_galore/) with the following setting: --clip_R1 4 --clip_R2 4 --three_prime_clip_R1 4 --three_prime_clip_R2 4. Trimmed read pairs were mapped to the reference genome (mouse: mm10; human: hg38) using Bowtie2 (11) with the following arguments: --dovetail --no-unal --no-mixed --no-discordant --very-sensitive -I 0 -X 1000. Prior to peak calling PCR duplicates and mitochondrial reads were removed and only properly aligned read pairs were used for peak calling with MACS2 (12) using ‘macs2 callpeak -f BAMPE -m 4 100 --bdg -SPMR’.

In order to compare NicE-seq data generated from different protocols, different numbers of input cells, different samples or compare NicE-seq to other open-chromatin mapping methods (i.e., ATAC-seq, DNase-seq), we downsized the mapped reads from different experiments to the same number of mapped fragments (after excluding PCR duplicates and mitochondrial reads) through random sampling. Peaks were called using the same parameter with MACS2 as mentioned above. Bigwig files of normalized reads per million read in 10 bp non-overlapping windows across the genome were displayed in Integrated Genomics Viewer (13). All analysis were performed after removing ENCODE blacklists.

#### Fraction of reads in peaks (FRiP)

The FRiP score was calculated using the deepTools plotEnrichment function (14). Called peaks were classified into 2 groups: TSS peaks if they overlap with +/−500bp from annotated TSSs (based NCBI RefGene annotation); and distal peaks if otherwise. Correspondingly, reads that overlapped with the TSS peaks by at least 1 base were marked as “TSS” reads. Reads that overlapped with distal peaks were marked as “distal” reads. Reads that do not overlap with any called peaks were marked as “reads not in peaks”.

#### Peak overlap analysis

Peaks called from different experiments were compared using the Bedtools (15). First peaks from all the samples are concatenated. Peaks that have at least one base pair overlapping are considered associated and are merged to form a union peak set. Then peaks of individual samples were compared to the union set and were marked as either “unique” or “common”. Last the numbers of “unique” and “common” peaks were summarized from all the samples and were used to make Venn Diagrams in R.

#### Correlation analysis

Correlation analysis of UniNicE-seq open-chromatin signals were performed with the DiffBind (16) package in R and deepTools (14) using two methods: occupancy (peak overlap) based method uses peak overlapping states and affinity (normalized read density) based method. The occupancy-based method determines the correlation coefficients based on the numbers of unique peaks and overlapping peaks. The affinity-based method first determines the number of normalized reads that overlap with a set of consensus peaks for individual samples and then calculates pearson correlation based on the normalized read count matrix. Correlation heat maps were generated using both occupancy and affinity methods with DiffBind and deepTools. PCA plots were generated from the normalized read count matrix by the affinity method.

#### Peak annotation and Gene/Genome Ontology analysis

Functional annotation of called peaks was performed with HOMER (17) annotatePeaks.pl. After associating peaks with nearby genes and assigning peaks to different genomic features (e.g., promoter, exon, CpG islands, repetitive elements etc), we also conducted Gene Ontology enrichment analysis for selected sets of UniNicE-seq peaks (e.g., tissue-specific peaks) and tested for enrichment of UniNicE-seq peaks in associated genomic features with HOMER.

#### Peak profile analysis in epigenetic relevant regions

To investigate enrichment of open-chromatin signals over sets of genomic regions with epigenetic significance, we first calculated normalized read coverage (number of reads normalized by the scaling factor of Reads Per Kilobase per Million mapped reads (RPKM)) in tiling bins of 100 bp across the entire genome from the bam file of properly aligned fragments (excluding PCR duplicates and mitochondrial reads) using the deepTools (11). Then heatmaps and profile plots were generated based on the normalized read coverage per 100 bp-bin over sets of genomic features of interest (TSS, enhancer, CTCF binding sites, RNA polymerase II binding sites) and the surrounding +/−2 Kb regions. As a control, the normalized coverage of a set of randomly sampled genomic regions were also plotted in the same way.

#### External datasets

TSS of mouse (mm10) and human (hg38) genomes were extracted from the NCBI RefGene gene table downloaded from the UCSC Table Browser. ChIP-seq datasets of cell-specific and tissue-specific CTCF binding, RNA polymerase II binding and histone marks (H3K27ac, H3K4me1, H3K4me3) were downloaded from the mouse and human Encyclopedia of DNA Elements (ENCODE) projects (Supp table 6). Human liver tissue-specific enhancers were acquired from the TiED database (http://lcbb.swjtu.edu.cn/TiED/) and EnhancerAtlas (18). The original hg19 genome coordinates were converted to hg38 using the LiftOver tool.

ATAC-seq (SRX2717891 & SRX2717892) and OmniATAC-seq (SRX2717893 & SRX2717894) datasets of mouse kidney were downloaded from NCBI SRA database. The DNase-seq dataset of mouse kidney (ENCSR000CNG) was acquired from the ENCODE project. The human HCT116 ATAC-seq and WGBS datasets were downloaded from NCBI GEO database (ATAC-seq: GSE101966 and WGBS: GSE97889). The DNase-seq dataset was acquired from ENCODE project (ENCSR000ENM).

## Data availability

All the UniNicE-seq data generated in this study is deposited in NCBI Gene Express Omnibus (GEO) under the accession GSE140276

## Acknowledgements

We thank W. Jack and C. Carlow for critical reading of the manuscript, T. Evans, D. Comb, Sir R.J. Roberts and J. Ellard for encouragement. Ninke Besbrugge for writing the stepwise detailed protocol. Basic research support for H.G.C., P.O.E, U.S.V. and S.P. was provided by New England Biolabs, Inc.

## Author contribution

H.G.C, P.C and P.O.E performed experiments. Z.S, U.S.V, C.H, and G.S performed data analysis. S.X purified enzyme and suggested experiments. H.W.L. designed and advised on experiments and analysis. S.P wrote the manuscript with input from all authors. S.P. supervised all aspects of this work.

## Supplementary Information

**Supp. Fig. 1.**
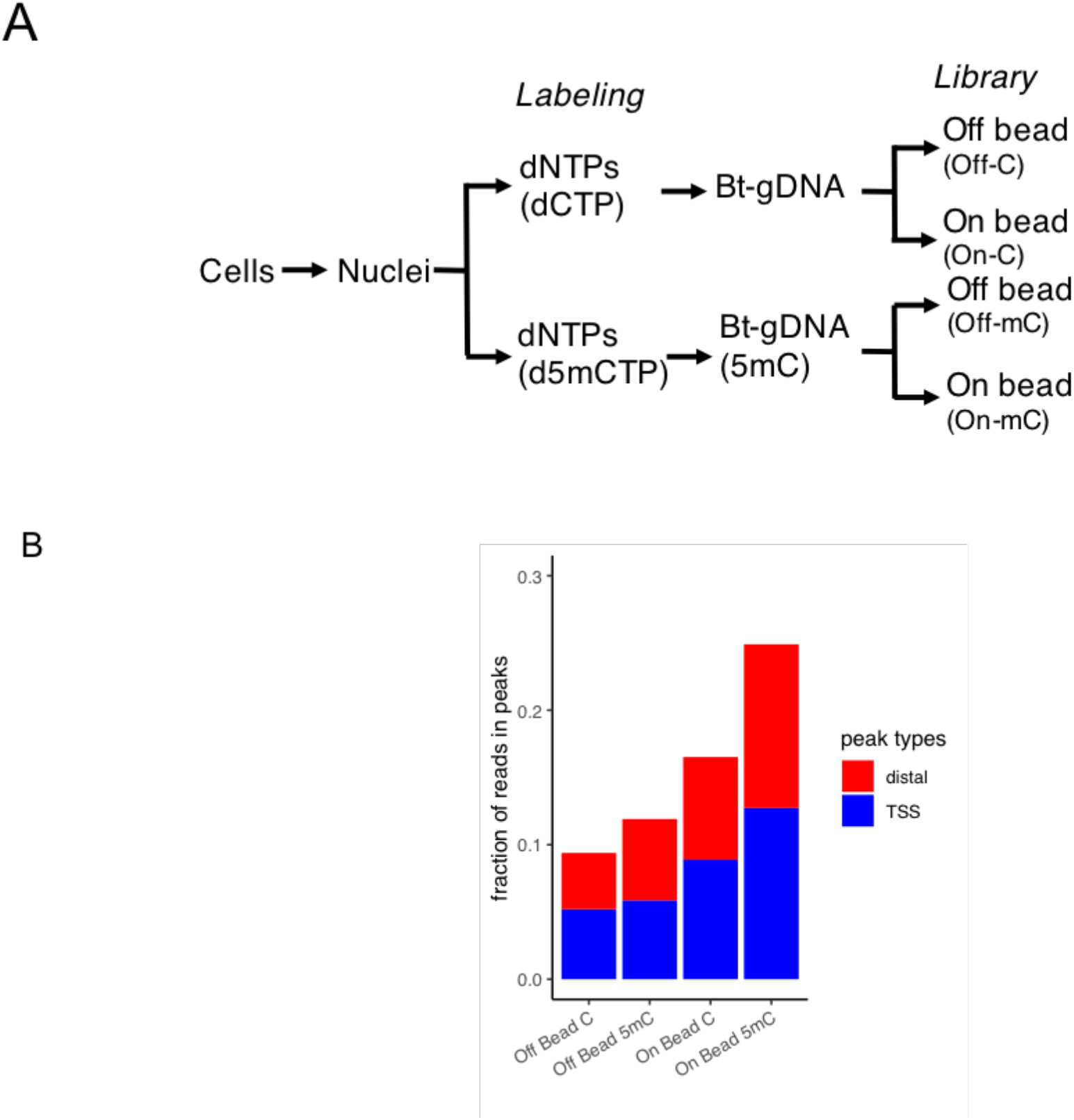
Optimization of universal NicE-seq (A) A schematic diagram of accessible chromatin labeling using dCTP or 5-mdCTP in the labeling reaction along with biotinylated-dCTP in the nucleotide mix. On-bead and off-bead represented presence of streptavidin magnetic beads for DNA capture and library preparation. (B) FRiP comparison between all 4 methods generated library that map to TSSs (+/−500bp of TSS) and distal elements (>500bp from TSS) from HCT116 cells. C and 5mC represents use of dCTP and 5-dCTP in the reaction mix.

**Supp. Fig. 2:**
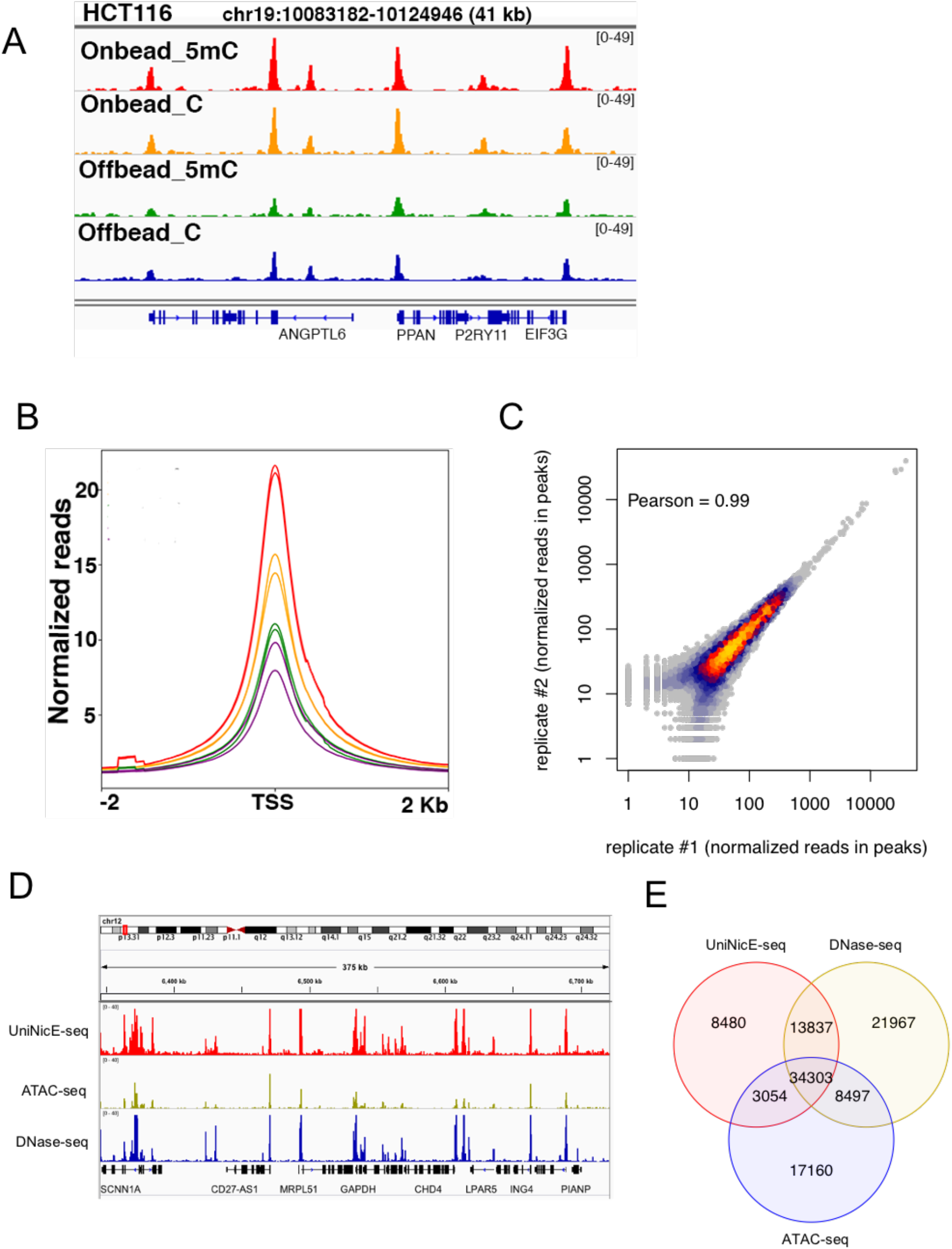
Optimization of accessible chromatin sequencing and comparison between UniNicE-seq, ATAC-seq and DNase-seq. (A) IGV screen shot of the normalized read density of the four NicE-seq conditions in HCT116 cells. (B) Distribution of the number of normalized HCT116 NicE-seq reads at transcription start sites (TSS) of human genes and the surrounding 2 Kb (− and +) regions. (C) Pearson correlation of normalized read densities in UniNicE-seq peaks of the 2 technical replicates in HCT116 demonstrating reproducibility. (D) IGV screen shot of the normalized read density of UniNicE-seq (top track), ATAC-seq (middle track) and DNase-seq (bottom track) in HCT116 (F) Overlap of HCT116 peaks called from 15M unique alignments using UniNicE-seq, ATAC-seq and DNase-seq.

**Supp. Fig. 3:**
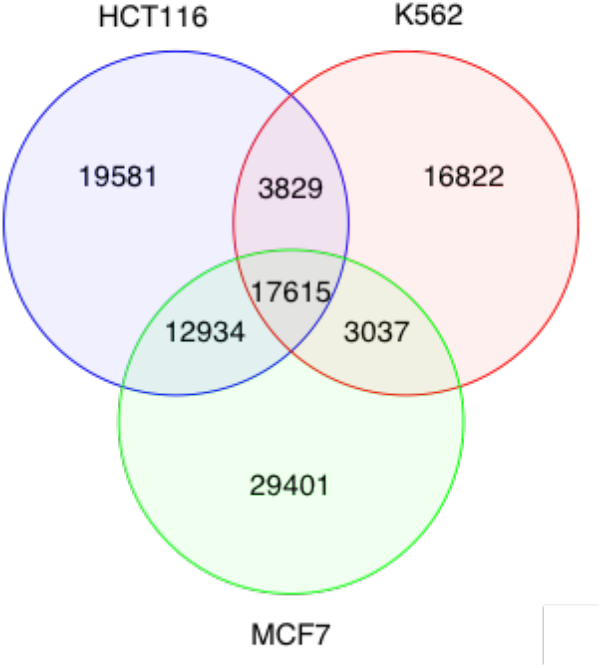
Venn Diagram showing common and cell-type specific UniNicE-seq peaks between the three cell types. (A) HCT116, K562 and MCF7 accessible chromatin regions were analyzed. Peaks are called from 11 million random sampled deduplicated alignment pairs.

**Supp. Fig. 4:**
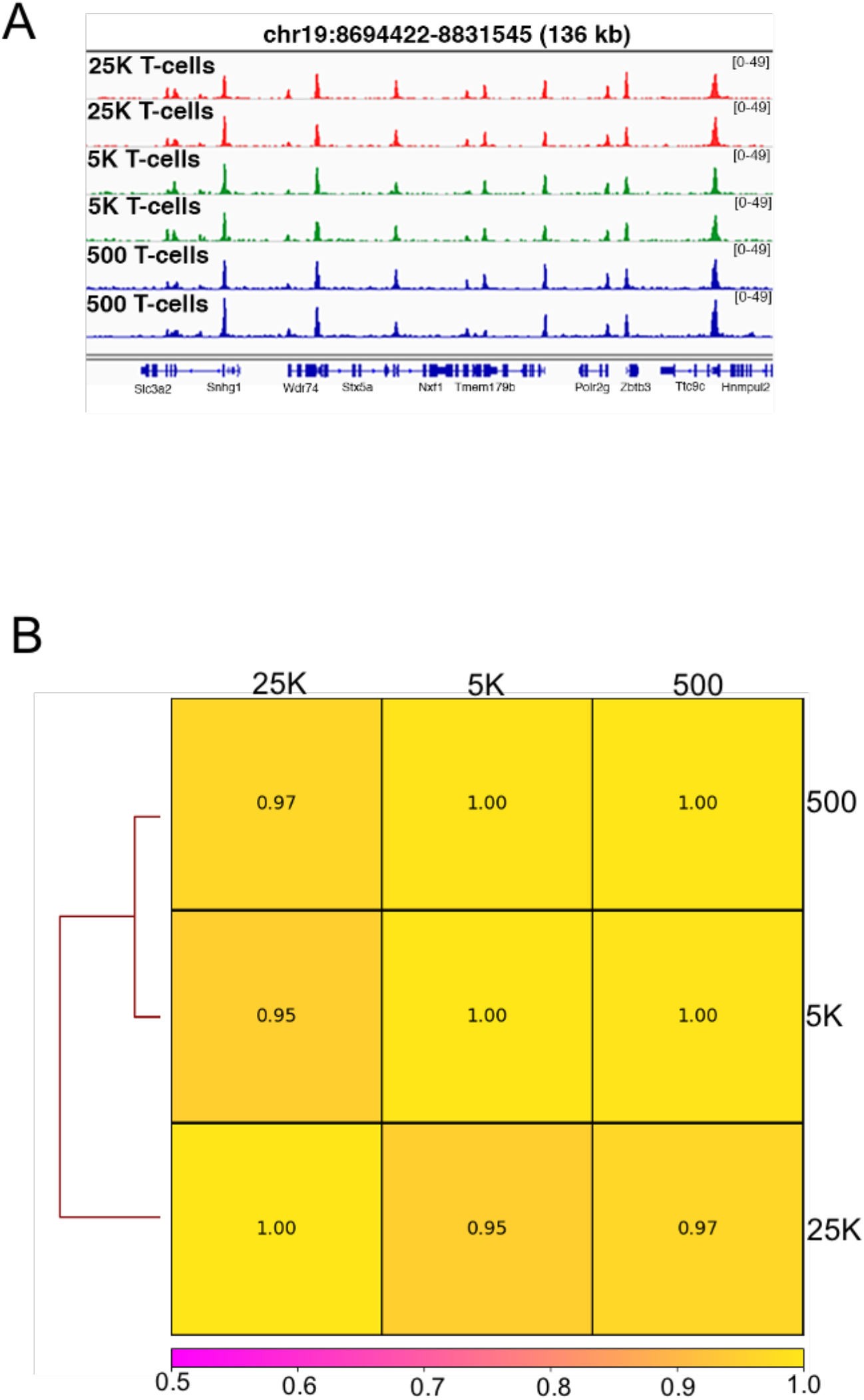
UniNicE-seq of mouse T cells cells. (A) IGV screen shot of the normalized read density of the technical duplicates of UniNicE-seq libraries of HCT116 cells at different cell numbers. (B) Pairwise comparison between all Universal NicE-seq reads between different T cell numbers from 500, 5 and 25K. Pearson’s correlation is indicated.

**Supp. Fig. 5:**
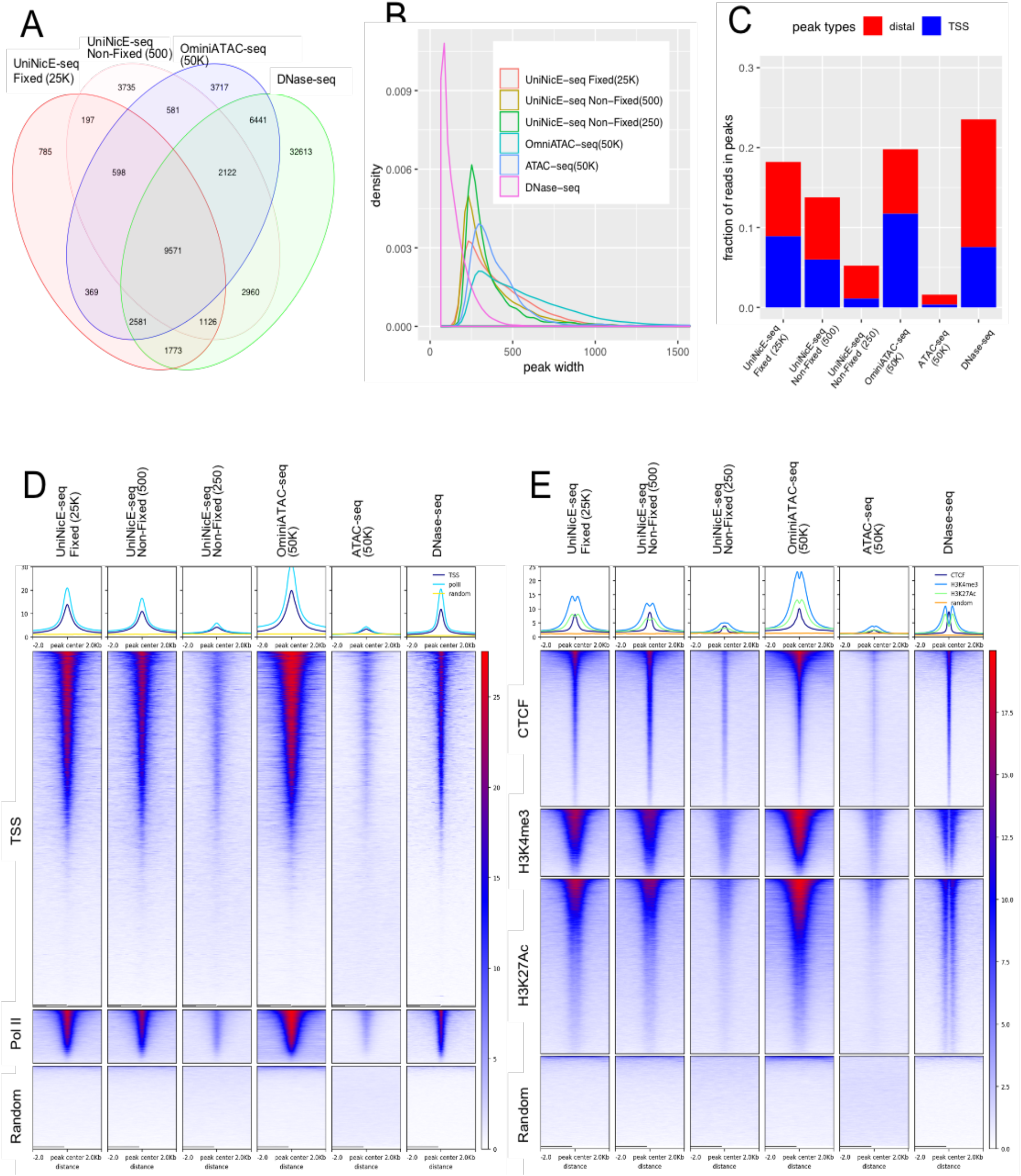
Comparison between UniNicE-seq, ATAC-seq, Omini ATAC-seq and DNase-seq of mouse kidney cells. (A) Venn diagram of accessible chromatin regions derived from UniNicE-seq, ATAC-seq, Omini ATAC-seq and DNase-seq of mouse kidney cells. (B) Distribution of fold change (FC) values (derived from MACS2) of the common accessible chromatin peaks of UniNicE-seq, ATAC-seq, Omni ATAC-seq and DNase-seq. (C) FRiP score of UniNicE-seq 25K, 0.5K and 0.25K compared with data obtained from ATAC-seq and OmniATAC-seq using 50K cells, and DNase-seq. (D) Heatmap showing comparison of normalized RPKM of 25K fixed, 0.5K nonfixed and 0.25 K mouse kidney UniNicE-seq, 50K omni ATAC-seq, 50K ATAC-seq and DNase-seq data at TSS, PolII and random. (E) Similar comparison like (D) along with chromatin features including CTCF, H3K4me3, H3K27Ac and random fragments.

**Supp. Fig. 6:**
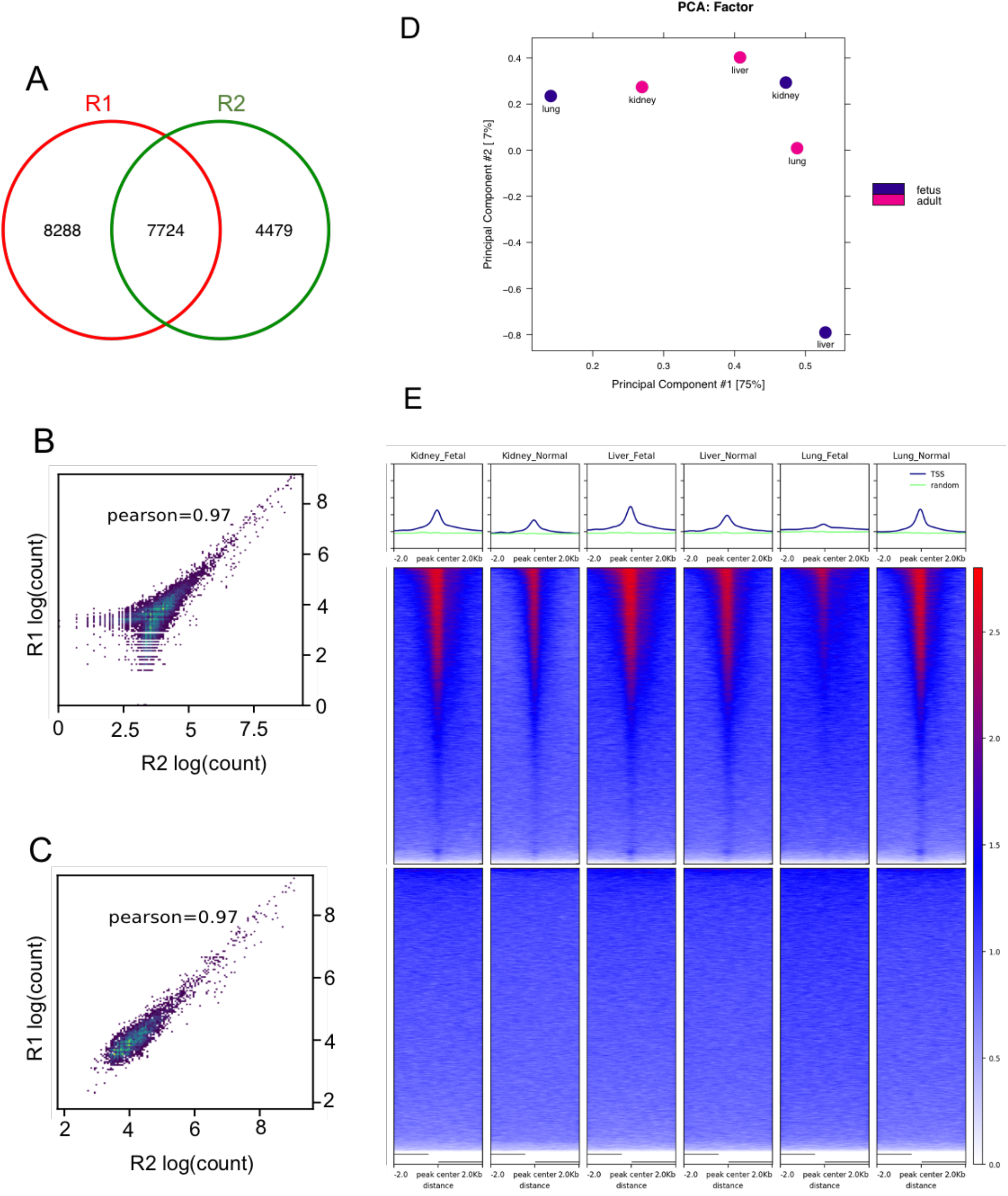
Comparison between accessible chromatin sequences two liver FFPE tissue section (A) Venn diagram demonstrating common accessible regions in two different human 5-10 μM lung normal tissue sections. (B) Pearson’s correlation analysis of total reads between two different human 5-10 μM lung normal tissue sections. (C) Pearson’s correlation analysis of common reads between two different human 5-10 μM lung normal tissue sections demonstrating quality of accessible peaks. (D) Principle Component Analysis and heat map of TSS across fetal and adult tissue for normalized read density of the consensus peaks between the samples from fetal and adult tissue. (E) Heat map of TSS (−/+ 2kb) between various tissue samples.

**Supp. Fig. 7:**
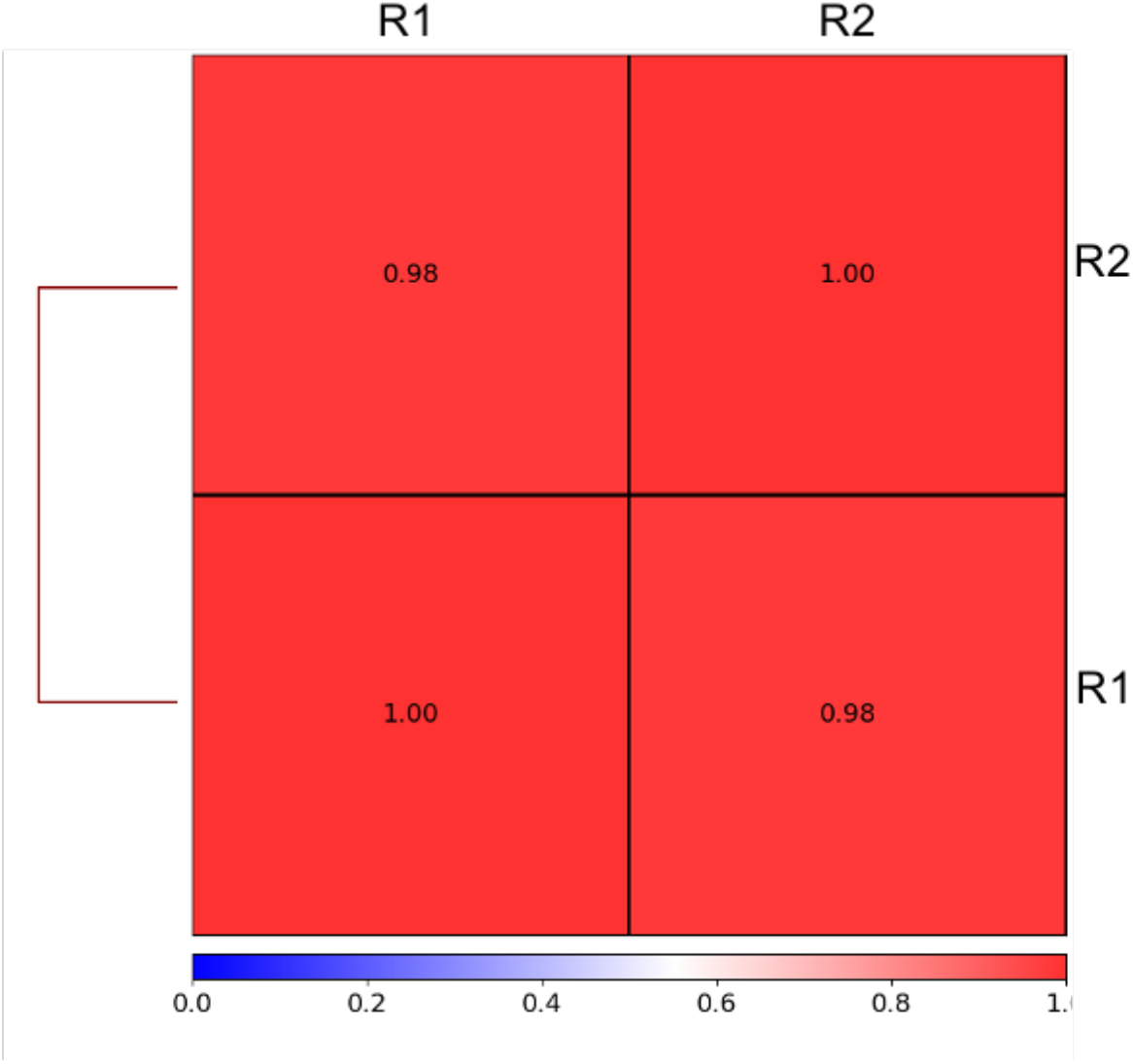
UniNicE-seq of normal human liver FFPE tissue section. Pearson’s correlation analysis by pairwise comparison between two different liver samples, R1 and R2.

**Supplementary Table 1:**
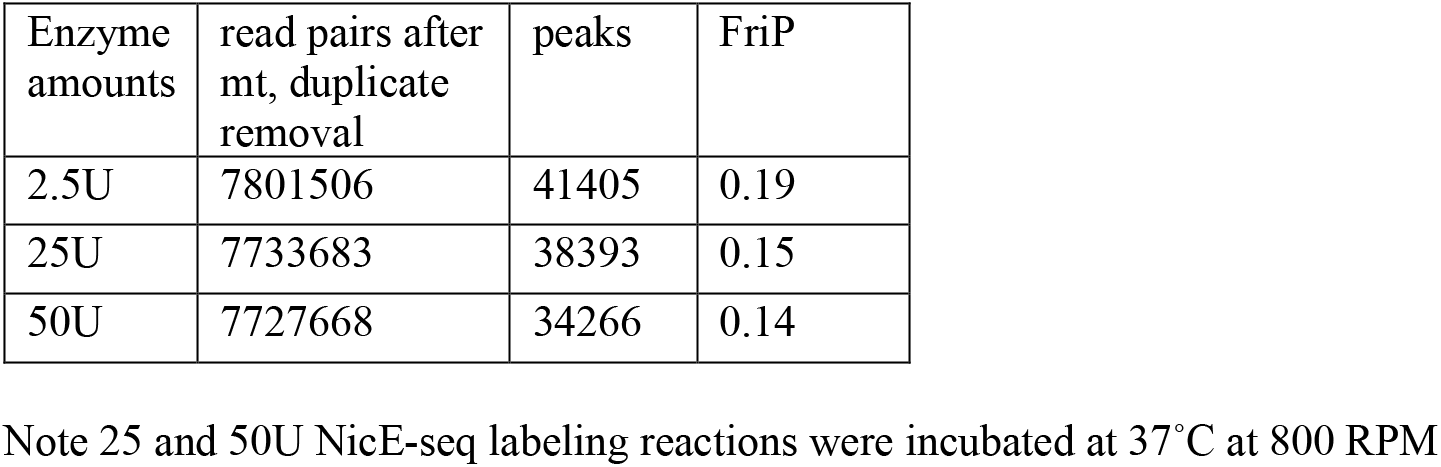
HCT116 UniNicE-Seq matrix for library with different amounts of enzyme.

| Enzyme amounts | read pairs after mt, duplicate removal | peaks | FriP |
| --- | --- | --- | --- |
| 2.5U | 7801506 | 41405 | 0.19 |
| 25U | 7733683 | 38393 | 0.15 |
| 50U | 7727668 | 34266 | 0.14 |

Note 25 and 50U NicE-seq labeling reactions were incubated at 37°C at 800 RPM

**Supplementary Table 2:** Quality control metrics of UniNicE-seq libraries applied to two human cell lines K562 and MCF7 in comparison to libraries made on and off beads with either with 5mdCTP or dCTP in the dNTP mix. We examined percentage of mitochondrial reads (“%mito”), number of total peaks and promoter peaks (+/− 500bp of TSS) and enrichment of signal at TSSs (“FRiP (TSS peaks)”). Two technical replicates were conducted for each sample. All the values were calculated from a subsample of 11 million de-duplicated alignment pairs.

| Cell Line | Bead | dCTP | replicates | %mito | total peaks | FRiP (all peaks) | TSS Peaks | FriP (TSS peaks) |
| --- | --- | --- | --- | --- | --- | --- | --- | --- |
| K562 | On | 5mC | 1 | 36.9 | 41723 | 0.18 | 12857 | 0.93 |
| K562 | On | 5mC | 2 | 33.9 | 37846 | 0.17 | 12456 | 0.87 |
| K562 | On | C | 1 | 15.1 | 28552 | 0.11 | 11385 | 0.64 |
| K562 | On | C | 2 | 17.7 | 29193 | 0.12 | 11437 | 0.66 |
| MCF7 | On | 5mC | 1 | 20.4 | 61703 | 0.22 | 14212 | 0.85 |
| MCF7 | On | 5mC | 2 | 22.1 | 63333 | 0.23 | 14284 | 0.89 |
| MCF7 | On | C | 1 | 12.1 | 47114 | 0.17 | 13310 | 0.72 |
| MCF7 | On | C | 2 | 14.7 | 54334 | 0.18 | 13747 | 0.77 |

**Supplementary Table 3.** Quality control metrics of UniNicE-seq libraries applied to mouse kidney tissues. 25K fixed cells were compared with 25K, 10K, 1K, 0.5K and 0.25K unfixed cells.

| Library | Raw read pairs | aligned read pairs | aligned% | Mito% | %dup | good alignment pairs | MACS2 narrow Peaks | FRiP |
| --- | --- | --- | --- | --- | --- | --- | --- | --- |
| nonfixed 250 | 40,654,273 | 39,449,830 | 98.3% | 0.19% | 63% | 14,198,157 | 11810 | 0.05 |
| nonfixed 500 | 30,509,137 | 22,963,486 | 92.1% | 2.60% | 59% | 9,019,473 | 30037 | 0.13 |
| nonfixed 5K | 33,665,133 | 32,557,812 | 97.9% | 1.29% | 17% | 26,653,887 | 59912 | 0.17 |
| nonfixed 10K | 39,287,749 | 38,255,104 | 98.5% | 0.39% | 10% | 33,903,582 | 51236 | 0.12 |
| nonfixed 25K | 39,736,105 | 38,675,476 | 98.4% | 0.04% | 6% | 35,867,394 | 57832 | 0.11 |
| Fixed 25K | 30,687,761 | 27,888,231 | 92.1% | 11.71% | 17% | 21,558,578 | 40716 | 0.18 |

**Supplementary Table 4a.** Quality control metrics of UniNicE-seq libraries applied to human adult lung tissues.

|  |  |  |  |  |  |  | 30M good alignment pairs |  |  |  |  |  |
| --- | --- | --- | --- | --- | --- | --- | --- | --- | --- | --- | --- | --- |
| Sample | Replicates | raw reads | alignment % | Mito % | duplication % | Good alignment pairs | Peak counts | FRiP in R1 peaks | FRiP in R2 peaks | TSS peaks | distal peaks | TSS% |
| Lung Adult 1 | 1 | 34990211 | 98.2 | 0.55 | 5.7 | 31322225 | 16024 | 2.6 | 2.1 | 5666 | 10358 | 35.4 |
| Lung Adult 2 | 2 | 39933602 | 98.6 | 0.51 | 3.6 | 36790808 | 12386 | 2.1 | 2.0 | 4110 | 8276 | 33.2 |

**Supplementary Table 4b.** Quality control metrics of UniNicE-seq libraries applied to different human adult and fetal tissues.^*^

| Sample | age | tissue | combined | narrowPeaks | TSS peaks | distal peaks | TSS peak% | FriP (all peaks) |
| --- | --- | --- | --- | --- | --- | --- | --- | --- |
| human tissue kidney fetal | fetus | kidney | R1+R2 50M alignment pairs | 38429 | 8411 | 30018 | 21.9% | 3.33 |
| human tissue kidney normal | adult | kidney | R1+R2 50M alignment pairs | 21500 | 4645 | 16855 | 21.6% | 2.48 |
| human tissue liver fetal | fetus | liver | R1+R2 50M alignment pairs | 35397 | 9248 | 26149 | 26.1% | 3.35 |
| human tissue liver normal | adult | liver | R1+R2 50M alignment pairs | 32890 | 6685 | 26205 | 20.3% | 2.98 |
| human tissue lung fetal | fetus | lung | R1+R2 50M alignment pairs | 16568 | 1091 | 15477 | 6.6% | 2.10 |

\*Here the replicates are merged together and downsized to 50M aligned pairs for the downstream analysis

**Supplementary Table 5.** Quality control metrics of UniNicE-seq libraries applied to human FFPE liver tissue sections.

|  | Replicates | Raw reads | Mapped reads | Alignment % | Mito % | Duplication % | Good aligned reads | Downsized to | Peaks | FRIP |
| --- | --- | --- | --- | --- | --- | --- | --- | --- | --- | --- |
| FFPE_Liver_1 | 1 | 117591288 | 93641034 | 79.63 | 5.70 | 24 | 49053108 | 49M | 50485 | 15.77 |
| FFPE_Liver_2 | 2 | 107390040 | 90729658 | 84.49 | 7.25 | 22 | 66428780 | 49M | 42816 | 14.55 |

**Supplementary Table 6.**
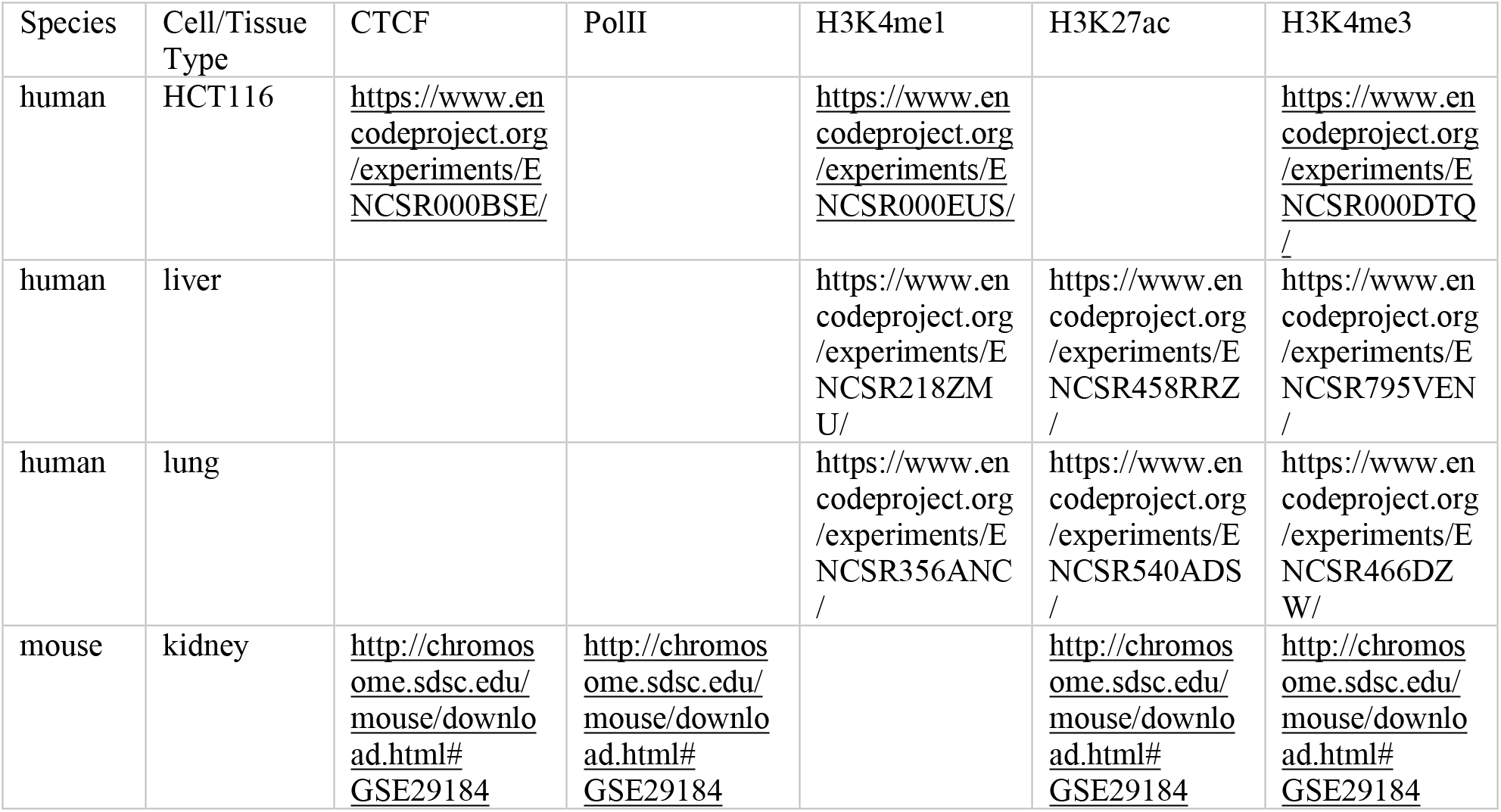
External ChIP-seq data sets of various human and mouse tissue and cell types in this work

